# Using serosurveys to optimize surveillance for zoonotic pathogens

**DOI:** 10.1101/2024.02.22.581274

**Authors:** E. Clancey, S.L. Nuismer, S.N. Seifert

## Abstract

Zoonotic pathogens pose a significant risk to human health, with spillover into human populations contributing to chronic disease, sporadic epidemics, and occasional pandemics. Despite the widely recognized burden of zoonotic spillover, our ability to identify which animal populations serve as primary reservoirs for these pathogens remains incomplete. This challenge is compounded when prevalence reaches detectable levels only at specific times of year. In these cases, statistical models designed to predict the timing of peak prevalence could guide field sampling for active infections. Thus, we develop a general model that leverages routinely collected serosurveillance data to optimize sampling for elusive pathogens by predicting time windows of peak prevalence. Using simulated data sets, we show that our methodology reliably identifies times when pathogen prevalence is expected to peak. Then, we demonstrate an implementation of our method using publicly available data from two putative ***Ebolavirus*** reservoirs, straw-colored fruit bats (***Eidolon helvum***) and hammer-headed bats (***Hypsignathus monstrosus***). We envision our method being used to guide the planning of field sampling to maximize the probability of detecting active infections, and in cases when longitudinal data is available, our method can also yield predictions for the times of year that are most likely to produce future spillover events. The generality and simplicity of our methodology make it broadly applicable to a wide range of putative reservoir species where seasonal patterns of birth lead to predictable, but potentially short-lived, pulses of pathogen prevalence.

**AUTHOR SUMMARY:** Many deadly pathogens, such as Ebola, Rabies, Lassa, and Nipah viruses, originate in wildlife and jump to human populations. When this occurs, human health is at risk. At the extreme, this can lead to pandemics such as the West African Ebola epidemic and the COVID-19 pandemic. Despite the widely recognized risk wildlife pathogens pose to humans, identifying host species that serve as primary reservoirs for many pathogens remains challenging. A key obstacle to confirming reservoir hosts is sampling animals with active infections. Often, disease prevalence fluctuates seasonally in wildlife populations and only reaches detectable levels at certain times of year. In these cases, statistical models designed to predict the timing of peak prevalence could guide efficient field sampling for active infections. Therefore, we have developed a general model that uses serological data to predict times of year when pathogen prevalence is likely to peak. We demonstrate with simulated data that our method produces reliable predictions, and then demonstrate an application of our method on two hypothesized reservoirs for Ebola virus, straw-colored fruit bats and hammer-headed bats. Our method can be broadly applied to a range of potential reservoir species where seasonal patterns of birth can lead to predictable pulses of peak pathogen prevalence. Overall, our method can guide future sampling of reservoir populations and can also be used to make predictions for times of year for which future outbreaks in human populations are most likely to occur.

## INTRODUCTION

Spillover of zoonotic pathogens is a pervasive challenge [1], imposing a persistent burden on human health and creating conditions ripe for the emergence of novel infectious disease [2]. One avenue to controlling the health impacts of spillover is to increase surveillance within the human population, treating disease as it occurs and using public health measures to keep initial events from expanding into epidemics or pandemics [3–5]. However, when surveillance and intervention systems fail, the results can be catastrophic (e.g., West African Ebola epidemic; COVID-19 pandemic).

An alternative approach to managing the risk of spillover is preemptive, and focuses on stopping spillover before it occurs. For instance, the risk of spillover could be managed by altering habitat availability for reservoir species [6, 7], changing human behavior to reduce contact with hosts [8, 9], or vaccinating reservoir species [10, 11]. For these preemptive approaches to work, we must know which animal species serve as important reservoirs for a pathogen of interest. Recent progress in this direction has been made by capitalizing on advances in machine learning that allow models to learn which suites of traits are associated with suitability as a reservoir [2, 12]. For instance, Schmidt et al. [13] used boosted regression trees to predict which species are most likely to serve as reservoirs for ebola viruses. Similar efforts have been used to predict reservoirs of SARS-CoV-2 [14], or- thopoxviruses [15], betacoronaviruses [12], Nipah virus [16], and filoviruses outside of equatorial Africa [17]. Thus, we now have tools in place to generate hypotheses for which species are likely to be reservoirs of any particular pathogen species.

Even with hypotheses for which species are likely to serve as a reservoirs in hand, testing and confirming that any individual species serves as an important reservoir remains a significant challenge [12, 18–20]. Beyond the obvious complexities and logistical challenges associated with sampling wild animals in remote locations, verifying that an animal is a reservoir requires capturing an animal with a detectable active infection [1]. Prevalence of some zoonotic pathogens is sufficiently high that screening reservoir animals for active shedding is straightforward (e.g., Lassa virus in *Mastomys natalensis* [21]), but more often it is extremely challenging for pathogens that generate short-lived acute infections concentrated at only certain times of the year [see 22–26]. In these cases, achieving even a modest chance of capturing an animal with a detectable active infection requires intensive and temporally focused sampling during periods of peak prevalence [18]. To address different aspects of this problem, several Bayesian approaches have been developed using serosurveillance data to predict incidence and prevalence in reservoir populations. For example, Borremans et al. [27] used information about multiple antibodies over time, pathogen presence, and demographic information to back-calculate the time since infection for individuals to estimate incidence of Morogoro virus infection in multimammate mice (*M. natalensis*). Using a different approach, Pleydell et al. [28] fit an age-structured epidemiological model specific to Ebola virus in straw-colored fruit bats (*Eidolon helvum*) to estimate the timing of peak prevalence in the adult population. Although these methods are robust, adapting them quickly to other systems would be laborious and not always feasible depending on data availability. Thus, a flexible method more easily tailored to different species that requires minimal data would aid empiricists developing surveillance sampling designs to target zoonotic pathogens.

Here we develop a general methodology that can be used to focus reservoir surveillance on periods of time that are most likely to coincide with peak prevalence of a zoonotic pathogen (most often viral pathogens). Our method requires routine serosurveillance data, knowledge of the rate at which detectable antibodies wane, and the rate at which individuals recover from infection. We validate our methodology using simulated data and then demonstrate an application by providing a workflow using R software [29] and previously published, free available serosurveillance data of Ebola virus (EBOV; *Zaire ebolavirus*) in straw-colored fruit bats (*Eidolon helvum*) and hammer-headed bats (*Hypsignathus monstrosus*). We believe this method is simple enough for wide-reaching application to many field studies that collect data on serology. Therefore our method provides a useful tool to guide the planning of field sampling and to study epidemiological dynamics in reservoir populations when data on active infections are rare or absent.

## METHODS

### Mathematical foundation

Our approach to optimizing surveillance for zoonotic pathogens from serological surveillance data builds from a mathematical model describing the ecology of the reservoir animal and the epidemiology of the pathogen. We illustrate our approach using a model of a reservoir animal that reproduces seasonally and experiences both density independent and density dependent mortality. We assume the pathogen can be adequately described by a modified SIR framework that takes into account both short-term antibody mediated immunity and long-term immunity mediated by a T-cell response. This distinction is important because we assume only the short-term antibody based response is detected by serology [30]. If we further assume individuals encounter one another at random, the ecology and epidemiology of the system can be described using the following system of differential equations:

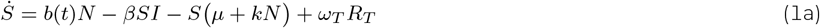

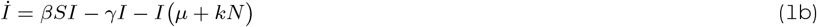

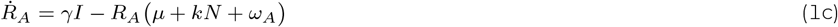

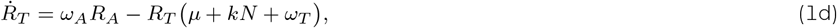

where *S* is the number of susceptible individuals, *I* is the number of pathogen infected individuals, *R*_*A*_ is the number of individuals with antibodies detectable through serology, *R*_*T*_ is the number of individuals that are immune to pathogen but lack detectable antibodies, and *N* = *S* + *I* + *R*_*A*_ + *R*_*T*_ is the total population size of the reservoir. All model parameters and their biological interpretations are described in table (1).

**Table 1:**
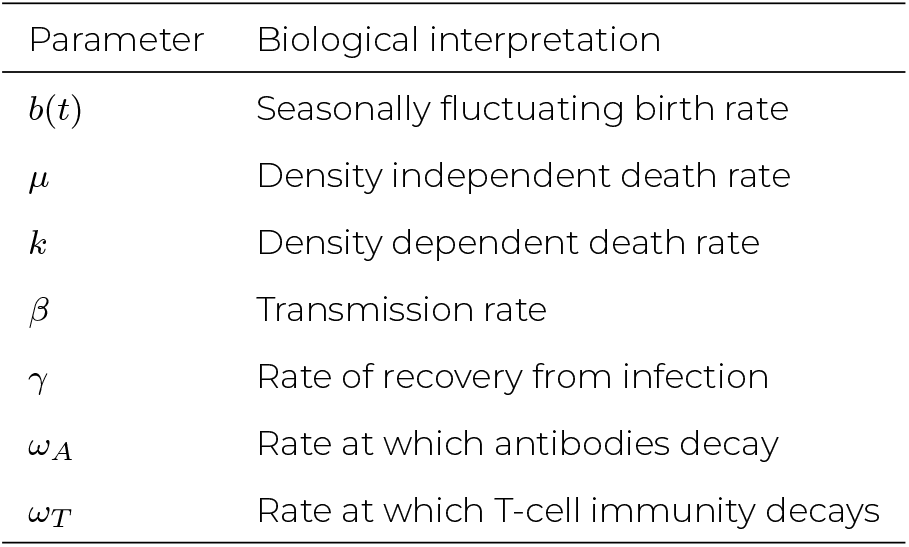
Model parameters and their biological interpretations. All rates are in days unless specified otherwise.

| Parameter | Biological interpretation |
| --- | --- |
| $b(t)$ | Seasonally fluctuating birth rate |
| $\mu$ | Density independent death rate |
| $k$ | Density dependent death rate |
| $\beta$ | Transmission rate |
| $\gamma$ | Rate of recovery from infection |
| $\omega_A$ | Rate at which antibodies decay |
| $\omega_T$ | Rate at which T-cell immunity decays |

If data on the abundance of each class are available, we could proceed directly from model (1). Unfortunately, this will not generally be the case, and data will more frequently come from serological testing of a random sample of *n* reservoir animals at various points in time. To calculate the probability that *x* animals will be seropositive within each sample of size *n* requires that we make a change of variables (supplemental material, appendix 1) to express model (1) in terms of proportions:

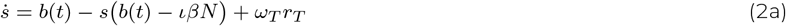

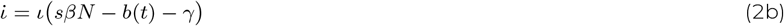

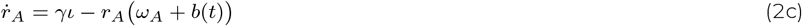

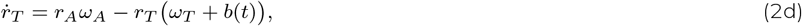

where *s, ι, r*_*A*_, and *r*_*T*_ are the proportion of reservoir animals in each class and *N* is the total population size of the reservoir animal. With the model now written in terms of proportions, we can proceed to solve for the proportion of animals in the actively infectious class, *ι*, as a function of the proportion of animals that carry antibodies, *r*_*A*_(*t*), using equation (2c):

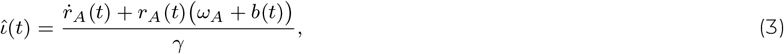

where 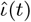 is the predicted proportion of the population that is actively infected at time *t*.

Equation (3) demonstrates that we can predict the proportion of the population that is actively infected at any point in time if we can estimate four quantities: 1) the rate at which antibodies are produced following infection, *γ*; 2) the rate at which antibodies wane over time, *ω*_*A*_; 3) a function describing the reservoir birth rate over time, *b*(*t*); and 4) a function describing seroprevalence over time, *r*_*A*_(*t*). We assume that the temporally constant parameters *γ* and *ω*_*A*_ are known or can be estimated using experimental infections in the lab. In contrast, the seasonal pattern of birth *b*(*t*) will generally not be known and may need to be estimated in some cases (supplemental material, appendix 2). If, however, animals live much longer than the lifespan of antibodies such that *b*(*t*) *<< ω*_*A*_, birth can be safely ignored to a good approximation (figure S1). Finally, we assume that the seasonal pattern of seroprevalence, *r*_*A*_(*t*), will generally be unknown and will need to be estimated from serosurveys. In the next section, we outline how this can be accomplished using routinely collected serological data. All mathematical analyses were performed in Wolfram Mathematica 13.1 [31].

### Fitting the mathematical model to data

Estimating a function that describes seasonal patterns of seroprevalence, 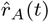, is central to our approach and leverages data that is routinely collected across a wide range of systems. In general, we assume a sample of reservoir animals is captured at multiple times each year and tested for the presence of antibodies for a target pathogen to give the number of seropositive animals in a sample. Thus, data will consist of a sampling date (*t*), a sample size (*n*), and the number of animals within the sample that are seropositive (*x*). We take two approaches to fitting 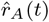, with the best approach dependent on the temporal resolution of the data and the users desire for credible intervals around the peak estimates.

#### Interpolation of temporally rich seroprevalence data

If high-resolution seroprevalence data (e.g., daily, weekly or monthly sampling) are available for a potential reservoir species, interpolation provides an efficient method for fitting the function *r*_*A*_(*t*) to the data. We illustrate this approach by applying a kernel smoother to estimate the function, 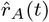. Specifically, we use the NadarayaWatson kernel regression estimate, with a normal density, available in R [29]. We then calculate the derivative of the interpolated function, 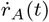, by differencing the fitted values for 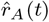 per unit time (e.g. days, weeks, months etc.). As long as parameters *γ* and *ω*_*A*_ have been estimated independently, and *b*(*t*) is negligible (or estimated), this provides the information required for the frequency of infected individuals over time, 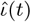 to be predicted using equation (3). Although computationally efficient and conceptually straightforward, interpolation only works within the range of the data and performs poorly when there are large gaps between data points. Thus, predictive performance using interpolation is expected to deteriorate when data are sparse or highly clustered (i.e., when sampling effort is concentrated at specific times of year).

#### Model fitting for sparse seroprevalence data

In cases when sampling is sporadic and seroprevalence data are sparse, interpolation may not be feasible and an approach based on model-fitting may perform better. This approach uses an understanding of system specific biology to define a mathematical function describing how seroprevalence is expected to change over time. The limited seroprevalence data is then used to estimate the parameters that fine-tune the function 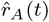 (e.g., the timing of peaks). Here, we illustrate this approach for systems where seasonal birth pulses are thought to cause fluctuations in the prevalence of infection and concomitant fluctuations in seroprevalence.

In systems where seasonal birth pulses occur, we expect, in general, a subsequent increase in infected individuals followed by a downstream increase in individuals that have seroconverted. Qualitatively, this expectation can be modeled using a modified periodic Gaussian function similar to [32]:

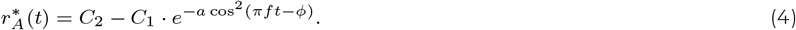

Here 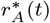 is a function specifying the predicted proportion of seropositive animals at time *t, C*_2_ adjusts the average value of seroprevalence over time, *C*_1_ sets the amplitude of seasonal fluctuations in seroprevalence, *a* controls the shape of seasonal fluctuations, *ϕ* defines the phase shift, and *f* specifies the frequency. We assume *f* is determined by the natural history of the reservoir species and is known. For example, a reservoir species that reproduces either once or twice per year in a regular pattern would have values of *f* = 1*/*365 and *f* = 2*/*365, respectively, if the time units are given in days. In contrast, we expect *C*_1_, *C*_2_, *a*, and *ϕ* to be unknown and require estimation. We emphasize 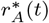 is an approximation to *r*_*A*_(*t*), which is unknown, designed specifically to predict the timing of the peaks. The result of using this approximation is the function 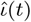 can take on negative values. Although negative values are biological not plausible, this approximation is still useful to recover the timing of the cycle.

We used Bayesian inference to estimate the unknown parameters in equation (4) and estimate the uncertainty in our estimates for 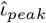 using 95% credible intervals (CI). Specifically, the likelihood of observing a temporal sequence of seroprevalence values is:

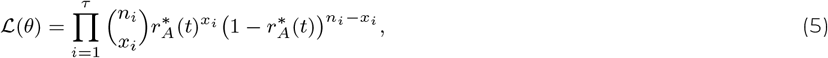

where the product is carried over *τ* total sampling time points, which occur over continuous time, *t*, and *θ* = *{C*_1_, *C*_2_, *a, ϕ}*^′^. For each sampling time point *i, n*_*i*_ defines the number of animals sampled and *x*_*i*_ defines the number of sampled animals found to be seropositive. Prior distributions for model parameters and details of the Bayesian estimation procedure are given in supplemental material, appendix 3. Bayesian estimation was performed using rstan [33].

### Simulating surveillance data

To determine if our methods accurately predict the true peak prevalence of infection, *ι*_*peak*_, we applied each method to simulated data sets. Specifically, we simulated a pathogen circulating in a wild animal population using model (1) with semi-annual birth pulses using equation (S6). In general, this leads to two peaks in prevalence and seroprevalence each year, a pattern observed in many bat species [e.g., 34–36]. Simulations focused on three different scenarios: low, medium, or high amplitude cycles in seroprevalence, *r*_*A*_(*t*), and prevalence, *ι*(*t*), with the specific parameter values used provided in table (S2). We generated 100 replicate stochastic simulations for each scenario using the Gillespie algorithm with a tau leaping approximation [37]. Simulations were initiated at the endemic disease equilibrium (supplemental material, appendix 1) and run for 10 years. We used the last 394 days for analyses to include peaks occurring at the end of year 9 to beginning of year 10, and all days in year 10. The two predicted peaks within the final 394 days, 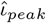, were determined for each simulated data set by finding the time point associated with the maximum value between days [0,170] and the time point associated with the maximum value between days [170,360].

We applied our methodology to the simulated data for a range of possible field sampling designs. First, we analyzed the simulated data sets assuming field sampling was performed at evenly spaced time intervals (daily, weekly, bi-weekly, monthly, bimonthly) over the 394 day study period. Second, we analyzed the simulated data sets assuming the number of sampling days was fixed at 42 days, but the distribution of these days over the year differed (evenly spaced days, random days, 3-day clusters, 7-day clusters). Each of the nine sampling designs was applied to the low, medium, and high amplitude seroprevalence cycle scenarios to yield 27 different combinations of epidemiological dynamics and sampling schemes (table S3). For each day of sampling, we assumed *n* = 20 animals were captured at random and tested for antibodies to the focal pathogen to yield an estimate for seroprevalence.

To evaluate the performance of our method, we compared the probability of detecting an actively infected animal (e.g., through PCR, culture, or sequencing) when sampling was timed using our method with three benchmarks: 1) the best case scenario where sampling was performed at the true peak, 2) a null hypothesis where sampling was performed on a random day, and 3) an alternative null hypothesis where sampling as performed 1*/γ* days prior to the peak in 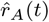. In each case, a sample of 20 animals was drawn at random and the number that were actively infected was recorded. Sampling was repeated ten times for each case and the probability of detecting an actively infected animal calculated as the number of trials in which at least one infected animal was found. Details on all simulations are given in supplemental material, appendix 4. All simulations were performed in R [29].

### Example study populations and surveillance data

To demonstrate an application, we apply our methodology to a previously published and freely available data set containing serosurveillance data for Ebolaviruses in two species of African fruit bats, straw-colored fruit bats (*Eidolon helvum*) and hammer-headed bats (*Hypsignathus monstrosus*). We use this data set to illustrate how our method can be implemented, how to meet the required assumptions, and the caveats arising when analyzing serosurveillance data.

*E. helvum* are commmon fruit bats that form large seasonal aggregations [38] and reproduce annually [39]. *H. monstrosus* also form large breeding aggregations [40], but unlike *E. helvum*, reproduce semi-annually [39]. Djomsi et al. [38] captured freeranging bats from a roosting site in Yaounde, Cameroon, and at a feeding site 40 km away near Obala, Cameroon. Samples were collected at approximately monthly intervals between December 2018 and November 2019, with the largest inter-sampling interval spanning two months. Whole blood samples and rectal and oral swabs preserved in RNA-later were collected from individual bats. Bat species, *E. helvum* and *H. monstrosus*, were identified by molecular testing. Djomsi et al. [38] screened *E. helvum* and *H. monstrosus* samples for antibodies to three *Ebolavirus* species using a Luminex-based serological assay previously adapted for bats [see 38]. They also tested for active infections in *E. helvum* using a semi-nested PCR assay specific to Ebola virus (EBOV; *Zaire ebolavirus*) targeting a 184 bp fragment on the VP35 gene [see 38]. For analyses in this study, we used the results from the Res1GP.ZEBVkiss antigenic test, a test for on the glycoprotein of EBOV, following [28]. Specific details of all methods and data are publicly available from [38] and [28].

The recovery rate and rate of waning antibodies to EBOV in *E. helvum* and *H. monstrosus* have not been estimated. Thus, we used measurements from experimental studies in Egyptian fruit bats (*Rousettus aegyptiacus*) with Marburg virus (MARV; *Marburg marburgvirus*) to approximate parameter values for the recovery rate (*γ* = 1*/*1.43 weeks) [41] and the rate of waning antibodies (*ω*_*A*_ = 1*/*12.9 weeks) [42]. We applied both methods, model fitting as the preferred method and then interpolation as a comparison, to each species in the example data set.

## RESULTS

### Optimizing surveillance on simulated data

#### Interpolation of temporally rich seroprevalence data

We begin our analyses by testing our methods on simulated surveillance data. Figure 1 shows an example of the true population curves for *r*_*A*_(*t*) and *ι*(*t*) and the estimated curves, 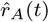 and 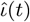, that were fitted to simulated serological data using interpolation. We find that we can successfully estimate 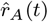 and predict prevalence pulses in populations with different epidemiological dynamics (e.g., low, medium, and high amplitude dynamics in figure 1) using interpolation. When surveillance sampling occurs at sufficient frequency and at even intervals across time, interpolation provides a good approximation to the true epidemiological dynamics such that surveillance sampling can be optimized to detect active infections (figures 2 and 3a). The accuracy of the predictions for the timing of peak prevalence in the population from interpolation for all sampling schemes are given in table (S4). When the data requirements are met, this method can also be used retrospectively to understand epidemiological dynamics when episodic shedding occurs randomly, for example, not necessarily coinciding with seasonal birth pulses (figure S7). However, interpolation methods can produce unreliable predictions for peak timing if serology data is sparse or sampling is highly clustered (see figure 3a). In these cases, model fitting is a better option because seasonal trends can be extrapolated between gaps in and outside of the range of the sampling data.

**Figure 1:**
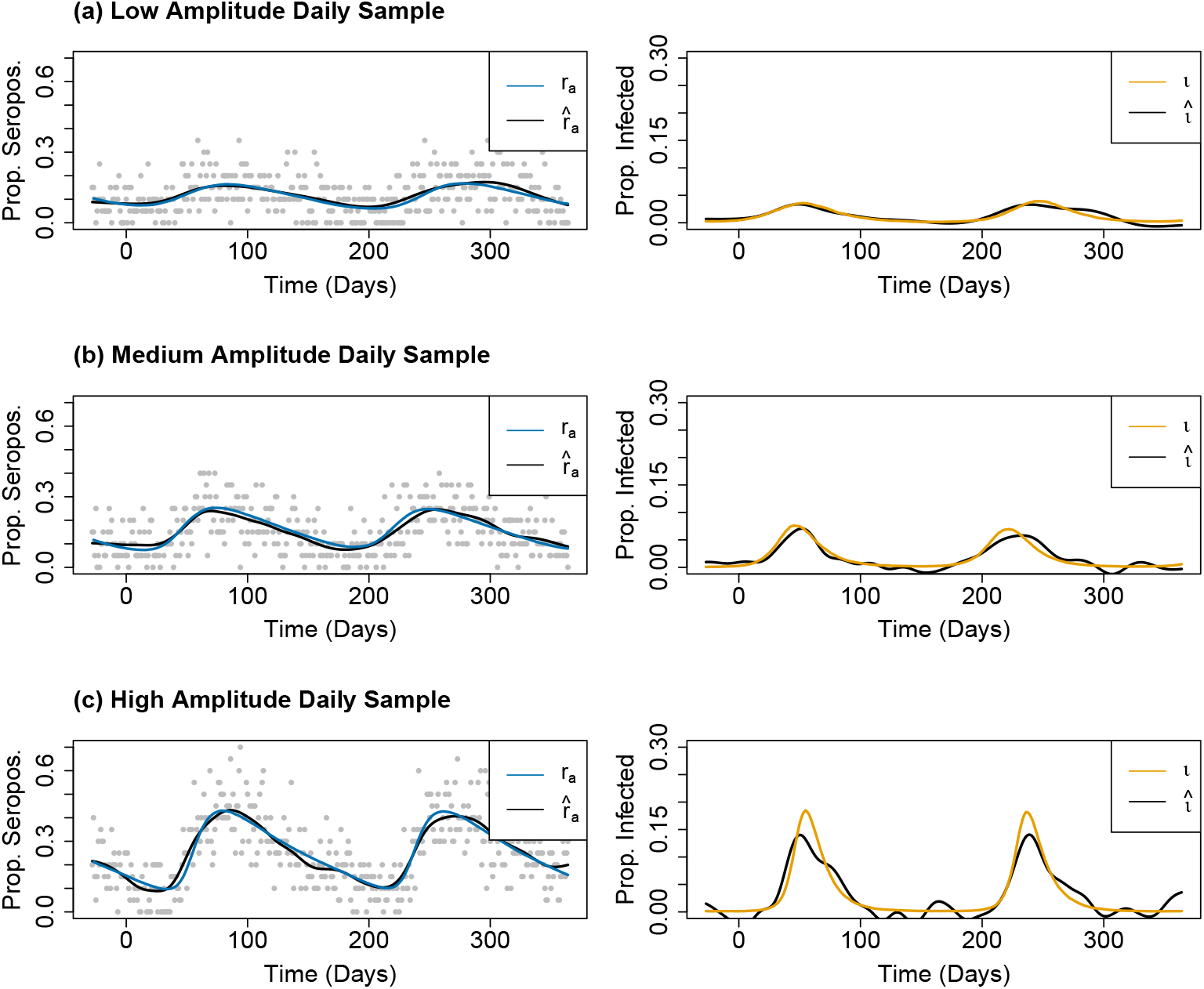
Results from interpolating serological data sampled daily for (a) low, (b) medium, and (c) high amplitude epidemic curves. Grey points represent the raw simulated data.

**Figure 2:**
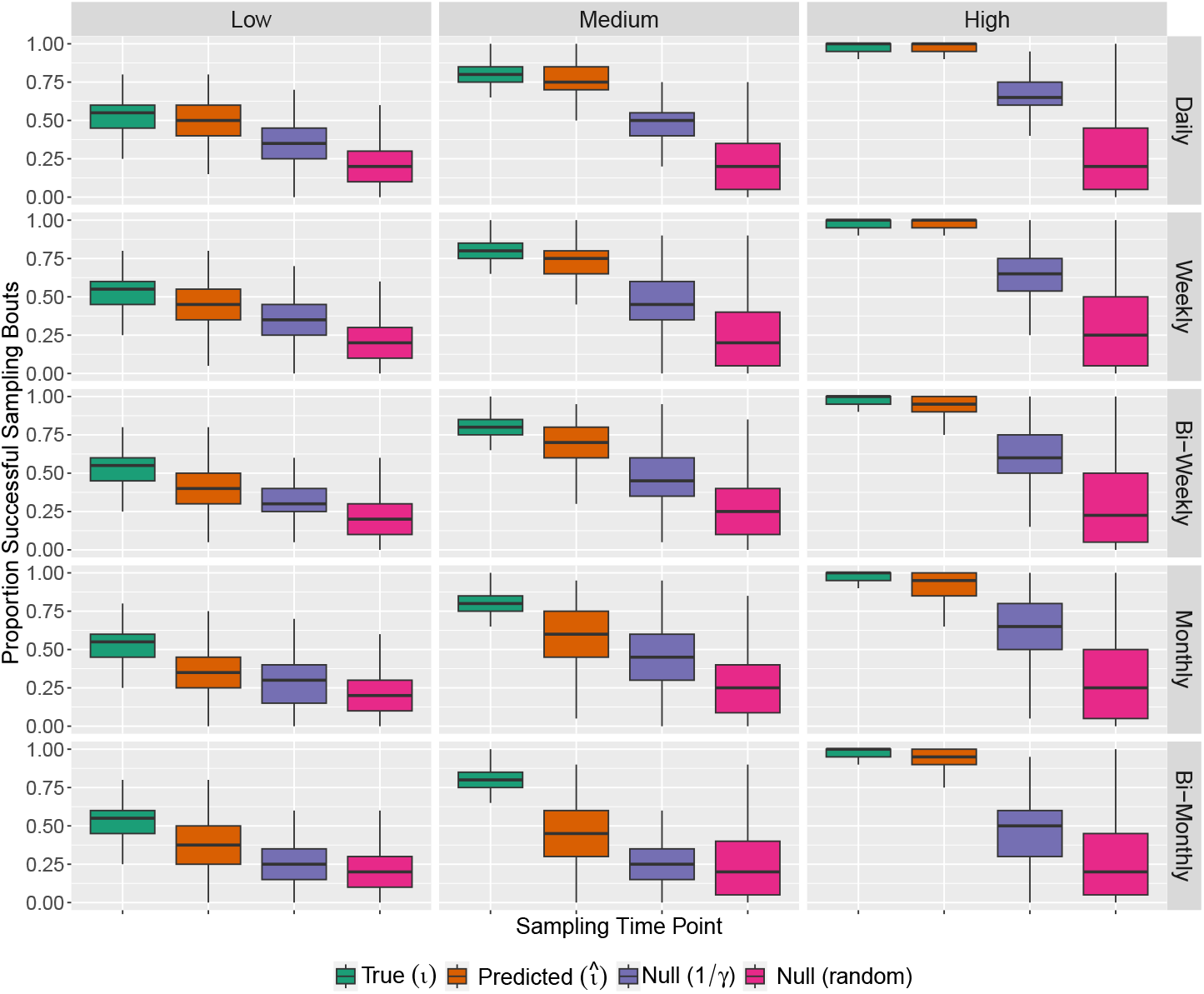
Proportion successful sampling bouts that occurred when the simulated population was sampled during the true peaks, *ι*_*peak*_, the interpolation prediction of peaks, 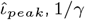 days prior to peaks of 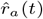, and a random time point. The proportion of successful sampling bouts are shown for three different types of disease dynamics, where the amplitude of the cycles is low, medium, and high, and for five different sampling schemes, when sampling occurs daily, weekly, bi-weekly, monthly, and bi-monthly over 394 days.

**Figure 3:**
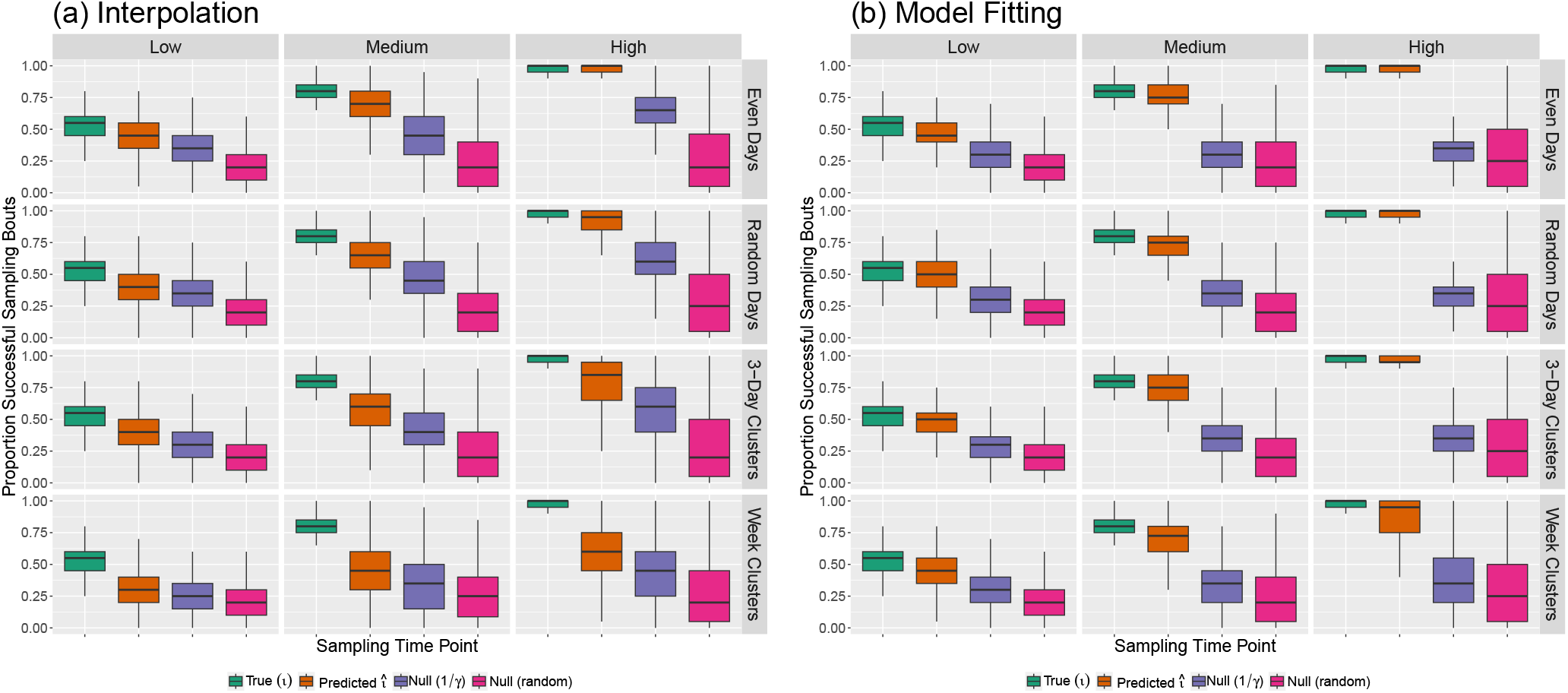
Proportion successful sampling bouts that occurred when the simulated population was sampled during the true peaks, *ι*_*peak*_, the predicted peaks, 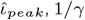 days prior to peaks of 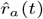, and a random time point when the predictions for 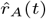 were made by (a) interpolation or (b) model fitting. The proportion of successful sampling bouts are shown for three different types of disease dynamics, where the amplitude of the cycles is low, medium, and high, and for four different 42-day sampling schemes, when sampling occurs at even intervals, random days, 3-day clusters, and week clusters.

### Model fitting for sparse seroprevalence data

Although estimating the function 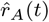 using model fitting is more computationally intensive than interpolation, our results show that this approach can accurately predict the timing of peak prevalence when interpolation is less reliable (figure 3). Fig-ure 4 shows an example of the true population curves for *r*_*A*_(*t*) and *ι*(*t*) and 500 estimated curves representing the credible intervals for 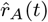 and 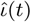. Specifically, as seroprevalence data becomes unevenly distributed, we find that the model fitting approach continues to provide accurate guidance for sampling whereas the guidance provided by the interpolation approach degrades (figures 3 and 4). The accuracy of the predictions for the timing of peak viral shedding in the population from 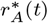, the size of the 95% CI estimated via Bayesian inference, and the proportion of times the true population peak falls within the CIs are summarized in table (S5). An added benefit of model fitting are the generation of credible intervals around the peak timing estimates.

**Figure 4:**
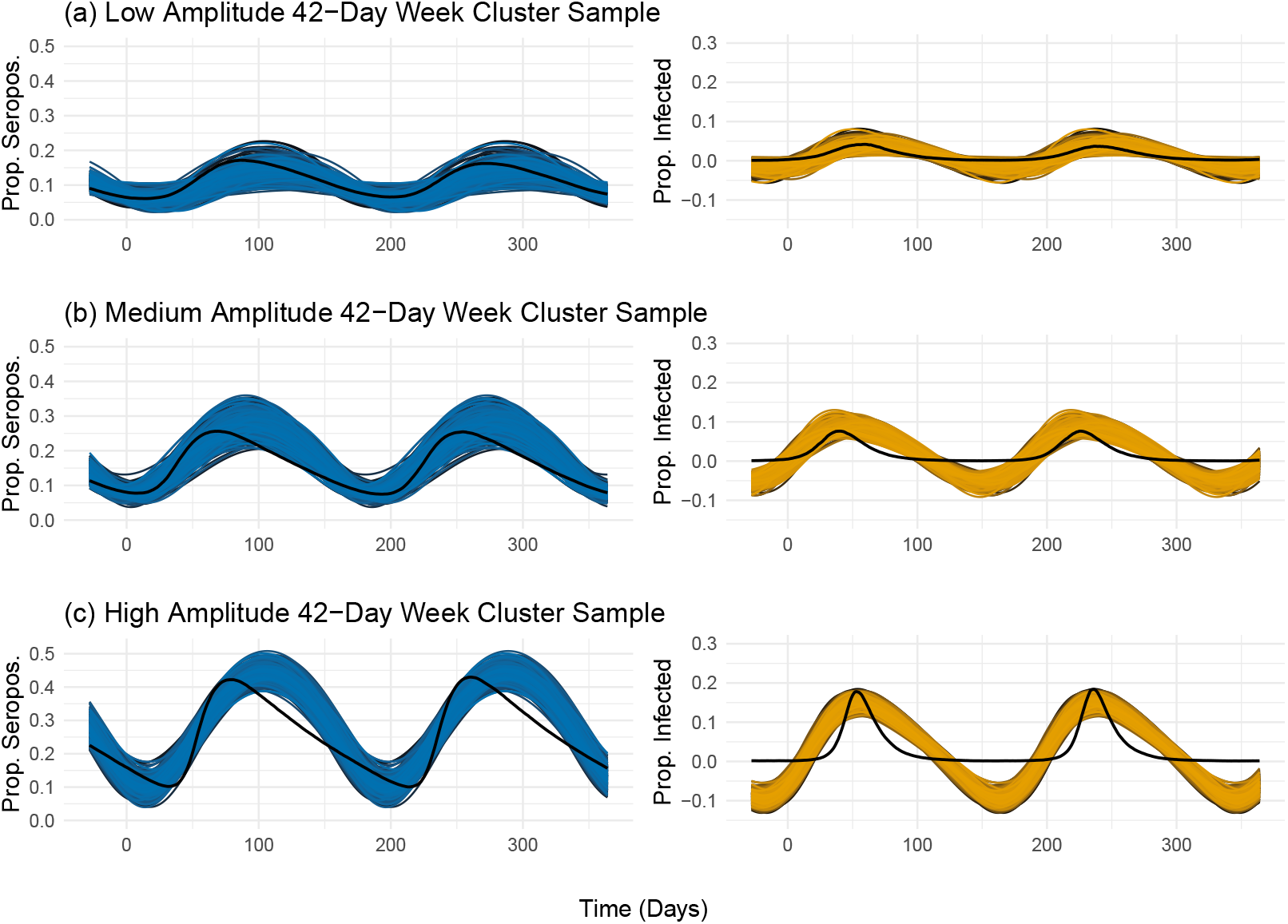
Results from model fitting with 42-day weekly clustered simulated serosurveillance data from (a) low, (b) medium, and (c) high amplitude epidemic curves. The blue and yellow lines represent the distribution of curves falling within the 95% CIs after Bayesian parameter estimation of 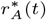 and predicting 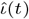, respectively. The black lines represent the true population simulated dynamics.

### Application to example data

We used publicly available data from [38] and [28] to predict the peak period, 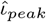, of active EBOV infections in *E. helvum* and *H. monstrosus* using our methodology. This data set includes seroprevalence and the proportion of animals lactating for each species over the period of one year. No active infections were detected in any *E. helvum* samples and *H. monstrosus* was not investigated for active infections [38]. Notable time gaps in sampling occurred from weeks 51 to 2 (weeks within a year are counted from from 0 to 51) and weeks 30 to 36. To begin our analyses, we tested the assumption that *E. helvum* reproduces annually and *H. monstrosus* reproduces semi-annually. Figures (S3) and (S5) demonstrate one annual birth pulse for *E. helvum* and two annual birth pulses for *H. monstrosus*, respectively.

Next, we used results from serosurveys to predict annual viral pulses, 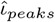, in *E. helvum* by fitting the data to 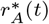 for EBOV and interpolating the data. Figure (5) shows the estimated serodynamics and predicted infection prevalence for this population from both methods, model fitting and interpolation. Results from model fitting suggest this population has a high amplitude cycle relative to our simulated data, with an average amplitude of 0.56 an 95% CI equal to [0.51, 0.62], meaning that sampling this population at peak prevalence greatly optimizes sampling for active infections. Our interpolation method predicted the peak to occur during week 34, and our model fitting method predicted the peak to occur during week 36 and 95% CI spanning weeks [35,37] (the distribution of the timing of predicted peaks is given in figure S4).

**Figure 5:**
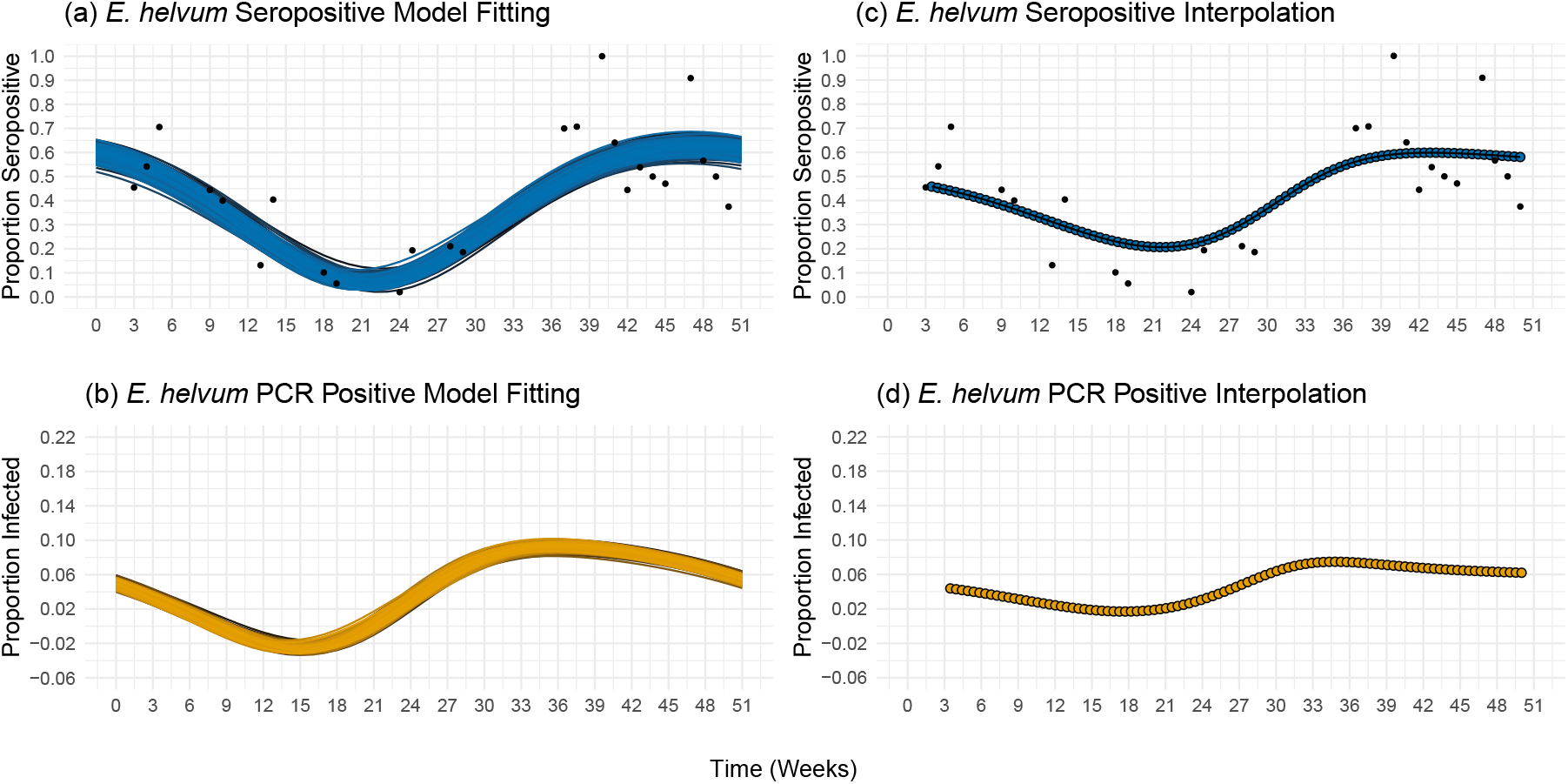
Predicted seasonal seroprevalence and prevalence patters in *E. helvum* for EBOV estimated from model fitting and interpolation. (a) Estimated seroprevalence curves from model fitting and sampling data (black dots) used to predict (b) prevalence over 52 weeks. (c) Estimated seroprevalence curves from interpolation and sampling data (black dots) used to predict (d) prevalence over 52 weeks.

We used the same methodology to predict the period of peak prevalence for *H. monstrosus*. The estimated temporal patterns of seroprevalence and prevalence for this population from both methods are shown in figure (6). Results from model fitting suggest that this population has a low amplitude cycle, with an average amplitude of 0.053 and a 95% CI equal to [0.00, 0.14]. The estimates and 95% CIs for the two peaks are the first mode occurring on week 27 within the interval [20.00, 32.00] and the second mode occurring on week 1 within the interval [46.00, 6.00] (the distribution of the timing of predicted peaks is given in figure S6). Using interpolation we can predict a weak peak occurring during week 30, but a second peak is not detectable. This could be due to only peak actually occurring the year the data was collected, or alternatively due to low the amplitude curves and because the timing of the peak occurs outside the range of the sampling data.

**Figure 6:**
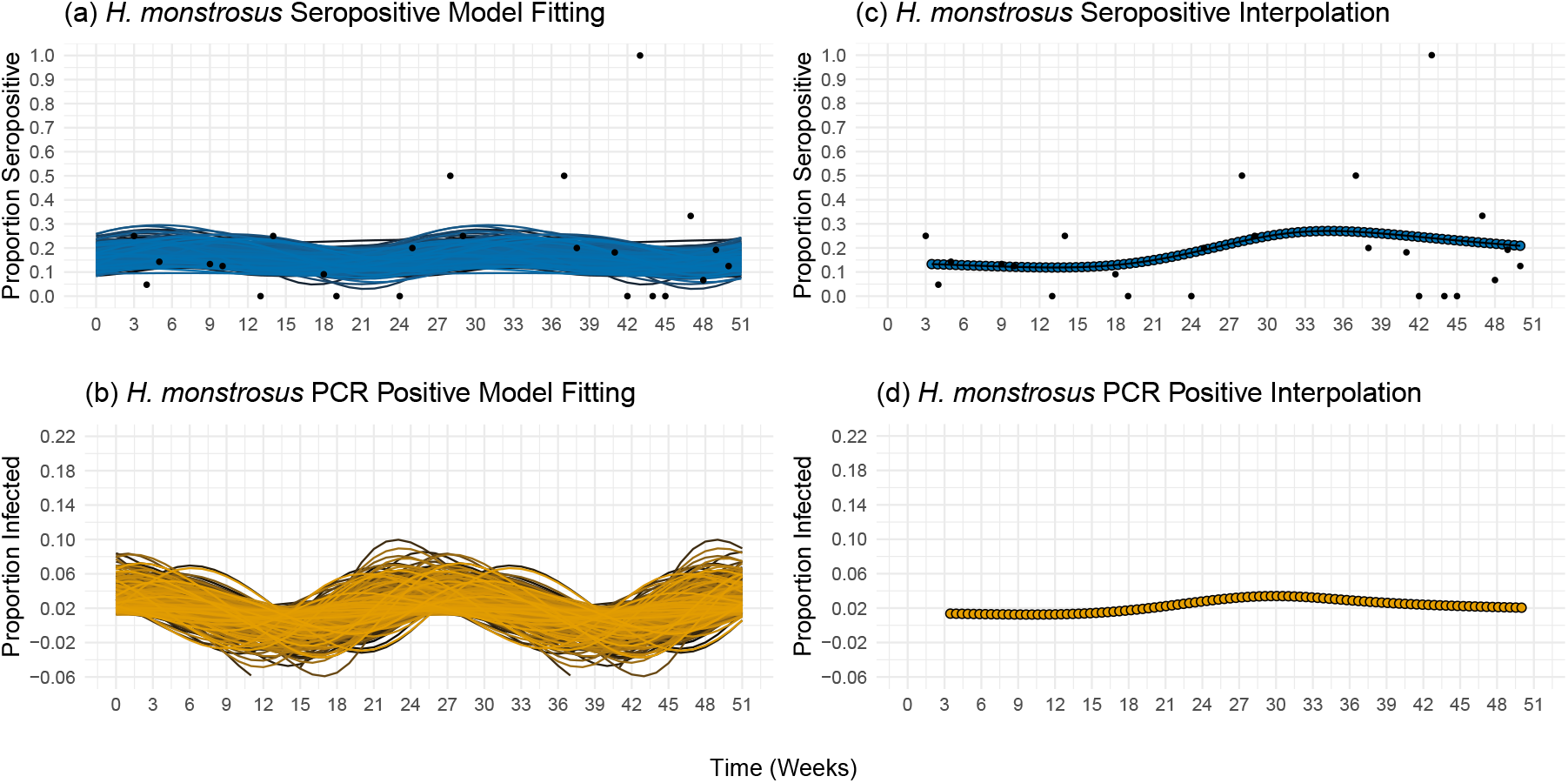
Predicted seasonal seroprevalence and prevalence patters in *H. monstrosus* for EBOV estimated from model fitting and interpolation. (a) Estimated seroprevalence curves from model fitting and sampling data (black dots) used to predict (b) prevalence over 52 weeks. (c) Estimated seroprevalence curves from interpolation and sampling data (black dots) used to predict (d) prevalence over 52 weeks.

## DISCUSSION

We have developed a general methodology for predicting the timing of peak pathogen prevalence in seasonally fluctuating wildlife populations using temporally structured serological data. Our approach is motivated by the possibility that successful sampling of actively infected reservoir animals has been impeded by seasonal fluctuations in pathogen prevalence driven by seasonal birth cycles. By focusing the search for active infections on specific periods of time where infections are most likely to be discovered, our method may facilitate confirmation of long-suspected reservoir hosts. Thus, our method leverages routinely collected serosurveillance data to extract information about the temporal pattern of active infection. When serosurveillance data is sufficiently rich such that the temporal pattern of seroprevalence to be interpolated, our method is particularly straightforward, computationally inexpensive, and accurate. Even when serosurveillance data is temporally sparse, our method can be used to generate accurate predictions by first fitting a mathematical model to the serological data. This latter approach, however, is more computationally intensive and requires an additional assumption about the timing of the birth cycle.

We apply our methodology to two bat species in Cameroon, *E. helvum* and *H. monstrosus* [38], hypothesized to harbor EBOV. We use this real-world example to illustrate an implementation of our methodology. Our results show that seasonal fluctuations in pathogen prevalence occur in these species. *E. helvum* had an annual prevalence peak and *H. monstrosus* had at least one peak occurring during the sampled year. Sampling at peak timing becomes more important as amplitude of the cycle increases (e.g., *E. helvum*), but even a small peak in prevalence (e.g., *H. monstrosus*) can still increase the probability of sampling an active infection. Therefore, seasonality should at least be considered before planning future field sampling. Validating our predictions in these species will only be possible, however, when animals with active infections have been captured from these populations. While unfortunate, this set back further highlights the knowledge gap that our methodology is contributing to filling.

Although the results from our simulated data are robust and empirical data demonstrate a real-world application, limitations of our model exist. First, the simulation testing assumed the mathematical model underlying our method accurately reflects the true biological processes. If the assumptions of our relatively simple compartment model are violated in the wild, our testing may overestimate the performance of our method. For instance, the model we have studied here ignores age structure which may have a significant impact on the relationship between seroprevalence and prevalence if sampling is not random with respect to age class [e.g., 28, 34]. The method we present here also assumes seasonality is driven by fluctuations in birth rate rather than seasonal changes in animal behavior that may influence contact rates and transmission [e.g., 40]. Even though we did not study these alternative scenarios directly, instead choosing to focus on a simple but general scenario, it will often be possible to integrate alternative biological assumptions by exchanging the underlying mechanistic model or estimating the frequency of the curve, *f*, along with the other unknown parameters if using model fitting.

Next, a potential limitation specific to our model fitting method is that we assume the epidemiological cycles occur consistently over time and the frequency can be specified using the number of birth pulses that occur annually for a particular species. In reality, prevalence pulses can occur stochastically [e.g., 43, 44], annual patterns in some population include skip years [e.g., 28, 45] or episodic shedding can be hard to distinguish from transient epidemics [46]. If epidemiological cycles cannot be approximated by a regular pattern, our model fitting method would not be appropriate. Our method also requires the rate of waning antibodies to either be known or estimated independently. Thus, our predicted peak intervals from model fitting are conditioned on specific values for rate of waning antibodies. If including uncertainty for these estimates is desired, our likelihood framework used in model fitting would easily accommodate a distribution for the rate of waning antibodies.

Last, our general method requires binary data describing whether an animal is seropositive or seronegative. Serological data is prone to cross-reactivity [47] resulting in low specificity and variable sensitivity dependent on the immune dynamics of the target species and pathogen, secondary antibody selection [48], and method of pathogen inactivation [49]. We assume reliable thresholds will be used to determine seropositivity, but we do not provide a method to include the uncertainty from serological data in our model. Caveats using serological data should be considered when interpreting the predictions.

Even in the face of these challenges and the limitations of our methodology, pathogen surveillance in wild animal populations is essential for identifying reservoir species, collecting pathogen samples for genetic characterization, and predicting when spillover is most likely to occur. By leveraging routinely collected serosurveys to optimize pathogen surveillance, the methodology we develop here has the potential to reduce the cost and labor associated with pathogen surveillance and increase our ability to successfully sample pathogens that reach appreciable prevalence at only specific times of year. More broadly, this simple methodology can be used to identify times of year when pathogen prevalence should peak, providing guidance for field sampling and interventions aimed at reducing spillover risk.

## ACKNOWLEDGMENTS

Funding was provided by the Centre for Research in Emerging Infectious Diseases East and Central Africa (CREID-ECA) grant number U01AI151799 in support of EC and SNS, PIPP Phase I: Predicting Emergence in Multidisciplinary Pandemic Tipping-points (PREEMPT) from the U.S. National Science Foundation (NSF) grant number 2200140 in support of EC, Verena (viralemergence.org) from NSF including NSF BII 2021909 and NSF BII 2213854 and the US National Institute of Allergy and Infectious Disease/National Institutes of Health (NIAID/NIH) grant number U01AI151799 in support of SNS, NIH 2R01GM122079-05A1 awarded to SNL and NSF DEB 2314616 awarded to SNL. Funding was also provided by the cooperative agreement CDCRFA-FT-23-0069 from the CDCs Center for Forecasting and Outbreak Analytics in support of EC and SNS. Its contents are solely the responsibility of the authors and do not necessarily represent the official views of the Centers for Disease Control and Prevention.

## DATA AVAILABILITY

All data used in this study was previously published and can be found online at at https://doi.org/10.5281/zenodo.8193102 from [28]. The data, R code and Mathematica Notebook used in this study can be found online at https://github.com/erinclancey/STOPPPModel.

