## Supplemental Material for "Using serosurveys to optimize surveillance for zoonotic pathogens"

### Supplemental material: Using serorsurveys to optimize surveillance for zoonotic pathogens

E. Clancey*[1], S.L. Nuismer[2] and S.N. Seifert[1]

[1]Paul G. Allen School for Global Health, Washington State University, Pullman, WA 99164 USA
[2]Department of Biological Sciences, University of Idaho, Moscow, ID 83844 USA

#### APPENDIX 1: SUPPLEMENT TO THE MATHEMATICAL FOUNDATION

Model (1) in the main text is a system of ordinary differential equations (ODEs) describing the ecology of a reservoir species and epidemiology of a pathogen expressed in counts. We use model (1) as the mathematical foundation of our method and to generate synthetic epidemics in a reservoir host. To find the pathogen free equilibrium for the population, $\mathcal{R}_0$, and the pathogen present equilibria for each disease compartment using model (1), we assume a constant birth rate by substituting the average birth rate $\bar{b}$ for $b(t)$. The total population size at equilibrium when the pathogen is absent using this assumption is

$$N_{eq} = \frac{\bar{b} - \mu}{k}. \tag{S1}$$

We can now use this pathogen free equilibrium to calculate $\mathcal{R}_0$ following the next generation method [1] giving us

$$\mathcal{R}_0 = \frac{\beta(\bar{b} - \mu)}{k(\bar{b} + \gamma)}. \tag{S2}$$

Using equation S1, we can also calculate the pathogen present equilibria for each compartment assuming birth rate is constant:

$$S_{eq} = \frac{\bar{b} + \gamma}{\beta} \tag{S3a}$$

$$I_{eq} = -\frac{(\omega_A + \bar{b})\left(\bar{b} + \omega_T\right)\left(\bar{b}(k - \beta) + \beta\mu + \gamma k\right)}{\beta k\left(\omega_A\left(\bar{b} + \gamma + \omega_T\right) + (\bar{b} + \gamma)\left(\bar{b} + \omega_T\right)\right)} \tag{S3b}$$

$$R_{Aeq} = -\frac{\gamma\left(\bar{b} + \omega_T\right)\left(\bar{b}(k - \beta) + \beta\mu + \gamma k\right)}{\beta k\left(\omega_A\left(\bar{b} + \gamma + \omega_T\right) + (\bar{b} + \gamma)\left(\bar{b} + \omega_T\right)\right)} \tag{S3c}$$

$$R_{Teq} = -\frac{\gamma\omega_A(\bar{b}(k - \beta) + \beta\mu + \gamma k)}{\beta k\left(\omega_A\left(\bar{b} + \gamma + \omega_T\right) + (\bar{b} + \gamma)\left(\bar{b} + \omega_T\right)\right)}. \tag{S3d}$$

We use the pathogen present equilibria as the initial values to simulate disease dynamics in our synthetic reservoir populations. Since we focus on a reservoir animal that experiences both density independent and density dependent mortality, we need to parameterize model (1) with a value for density dependent death rate, $k$, such that the population size reaches a stable equilibrium given birth rate, $b(t)$, and density independent death rate, $\mu$. To find $k$ that meets this criteria, we assume effective lifespan $(L)$ must be equal to $\frac{1}{b}$, and therefore

$$k = \frac{\frac{1}{L} - \mu}{N_{eq}} \tag{S4}$$

when the population is at equilibrium. Equation S4 was used to calculate the values for $k$ used for simulating surveillance data (table S2).

We assume serosurveillance data will most often be given in terms of proportion seropositive animals obtained within a sample. To convert model (1) to proportions we make a change of variables directly to the system of ODEs using the quotient rule to get model (2) as it appears in main text:

$$\dot{s} = \frac{\dot{S}}{\dot{N}} = \frac{N\dot{S} - \dot{N}S}{N^2} = b(t) - s\big(b(t) - \iota\beta N\big) + \omega_T r_T \tag{S5a}$$

$$\dot{\iota} = \frac{\dot{I}}{\dot{N}} = \frac{N\dot{I} - \dot{N}I}{N^2} = \iota\big(s\beta N - b(t) - \gamma\big) \tag{S5b}$$

$$\dot{r}_A = \frac{\dot{R}_A}{\dot{N}} = \frac{N\dot{R}_A - \dot{N}R_A}{N^2} = \gamma\iota - r_A\big(b(t) + \omega_A\big) \tag{S5c}$$

$$\dot{r}_T = \frac{\dot{R}_T}{\dot{N}} = \frac{N\dot{R}_T - \dot{N}R_T}{N^2} = r_A\omega_A - r_T b(t) - \omega_T r_T. \tag{S5d}$$

#### APPENDIX 2: SUPPLEMENT FOR THE ESTIMATION OF SEASONAL BIRTH PULSES

If the information is available and adding information about fluctuating birth pulses to the model is needed, below we offer a method to estimate $b(t)$. Similar to serosurveillance data, we assume a sample of reservoir animals is captured at multiple times each year. The data that can be used to estimate seasonal patterns of reproduction are diverse and may include, for example, the proportion of pregnant females, the proportion of juvenile animals, or the proportion of lactating females over time. Thus, data will consist of a discrete sampling date ($i$), a sample size ($n_i$), and the number of animals within the sample that were birthed (or a metric thereof) ($x_i$). Given any of these data examples, we use the cyclic Gaussian function described and tested by [2]:

$$b(t) = g \cdot e^{-s\cos^2(\pi f t - \psi)}, \tag{S6}$$

where $g$ controls the amplitude, $s$ controls the shape of the cycle, $f$ is the frequency, $\psi$ is the phase shift, and $t$ is continuous time.

However, we show the effect of seasonal birth rate, $b(t)$ in equation (S6), on $\hat{\iota}$ is negligible as long as $b(t) << \omega_A$. This occurs because $b(t)$ and $\omega_A$ appear only as a sum in equation (3) in the main text. When birth rates are an order of magnitude or more smaller than $\omega_A$, $b(t) + \omega_A \approx \omega_A$. We show this graphically in figure (S1) using the numerical solution for $\iota$ from equation (2b) and $\hat{\iota}$ from equation (3) in the main text.

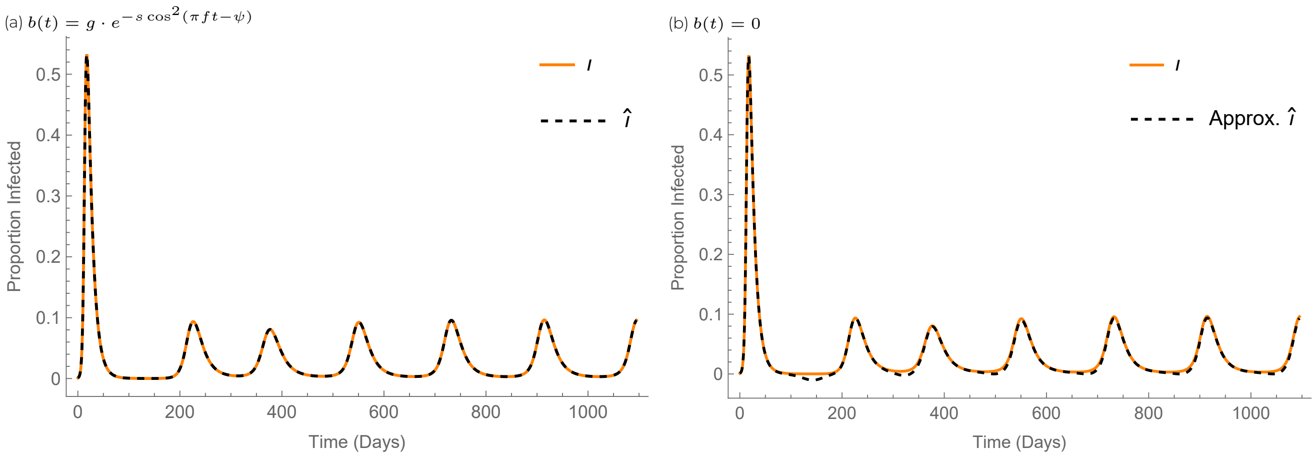


**Figure S1:** We compare the true numerical solution for $\iota$ from model (2) in the main text to $\hat{\imath}$ from equation (3) in the main text with (a) $b(t)$ included in the equation and (b) $b(t)$ set to zero.

Nonetheless, to fit equation (S6) to data, we must estimate the unknown parameters $g$, $s$, and $\psi$. We assume $f$ to be a known parameter based on natural history of a species reproductive cycle with the same value as in equation (4) in the main text. Since we assume reproductive data will also be in proportions, we can use equation (5) in the main text by exchanging $b(t)$ for $r_A^*(t)$ to give the probability of a successful trial and with $\theta = \{g, s, \psi\}'$.

A noticeable problem arises when fitting data on proportion captured animals that are pregnant or lactating instead of data on birth rates directly. When the estimation is performed on, for example, proportion lactating females, this specifies a function that is proportional to $b(t)$, preserving the shape and timing, but not the amplitude in units of per capita birth rates. Once the parameters are fitted, this problem can be rectified because we can adjust the value of $g$ if we know or can estimate the average birth rate of the population $\bar{b}$ with the following equation:

$$g = \frac{\bar{b} \cdot \frac{1}{f}}{\int_0^{\frac{1}{f}} e^{-s\cos^2(\pi f t - \psi)} dt}. \tag{S7}$$

Now, with the estimates for $\theta$ with adjusted $g$, the parameterized equation (S6) can be plugged into equation (3) in the main text to calculate $\hat{\imath}$ if this is needed.

#### APPENDIX 3: SUPPLEMENT TO THE BAYESIAN ESTIMATION

We take a Bayesian approach to parameter estimation to both maximizing the likelihood function in equation (5) and quantifying the uncertainty in our estimates for $\hat{\imath}$. We used the rstan package which implements the Hamiltonian Monte Carlo algorithm [3]. We allowed the algorithm to run for 1,000 iterations for the simulated data and 10,000 iterations to obtain estimates on the empirical data. For each serology dataset, simulated or empirical, we re-sampled each marginal posterior distribution with replacement for $n = 500$ and then kept all parameter values falling within the 95% credible intervals (CI). All credible intervals were calculated as highest density intervals (HDI) using bayestestR [4]. The remaining values in the joint posterior distribution were then used to generate the curves in figures

(4,S4,S6) and estimate the predicted peaks for the simulated and empirical data. Details on the prior distributions for all unknown parameters in equation (4) in the main text and equation (S6) are given in table (S1) (equation (S6) was used to fit the curves in figures S3 and S5). All prior distributions were uniform and bounded the posterior distributions such that the resulting serology curve remained within the interval $[0, 1]$ to model proportions, produced a biologically plausible serology curve and derivative, and only allowed estimation to occur over a single period.

**Table S1:** Prior distributions use for Bayesian inference. All priors are uniform with the same bounds for rates in days or weeks.

| Parameter | Equation | Prior Distribution |
|---|---|---|
| $C_1$ | 2.4 | $\mathcal{U}[0, 1]$ |
| $C_2$ | 2.4 | $\mathcal{U}[C_1, \frac{1+C_1}{e^a}]$ |
| $a$ | 2.4 | $\mathcal{U}[0, 1]$ |
| $\phi$ | S6 | $\mathcal{U}[0, \pi], [\frac{-\pi}{2}, \frac{\pi}{2}]$ |
| $g$ | S6 | $\mathcal{U}[0, 1]$ |
| $s$ | S6 | $\mathcal{U}[0, \infty]$ |
| $\psi$ | S6 | $\mathcal{U}[0, \pi], [\frac{-\pi}{2}, \frac{\pi}{2}]$ |

#### APPENDIX 4: DETAILS ON THE SIMULATED SURVEILLANCE DATA

We generated 300 synthetic reservoir populations with seasonal forcing of a circulating pathogen over a period of 10 years. We engineered 100 low, medium, and high amplitude epidemic curves to use different scenarios to test the ability of each method to predict $\hat{\imath}$. We simulated each population using model (1), but also added migration to get more consistent epidemics (failed epidemics reduces computational efficieny). Datasets were removed if prevalence or amplitude were zero, and we kept the last 394 days for analysis. Amplitude was calculated as the difference between the peak and the trough (note this differs from a standard amplitude calculating which is the difference between the peak and the midline). We manipulated amplitude by changing the birth and death rates of the reservoir population. Exact parameter values for population simulations are given in table (S2), an example of the low, medium, and high amplitude cycles are shown in figure (S2), and population information on amplitude, seroprevalence, prevalence, $\mathcal{R}_0$ are given in table (S3).

**Table S2:** Parameter values used to generate the 100 low, medium, and high amplitude simulated epidemics. When parameter values vary, they correspond to low, medium and high epidemic cycles in that order. All rates are given in days.

| Parameter | Definition/Reference | Value in Simulations |
|---|---|---|
| $N_0$ | Initial pop. size | $1 \times 10^5$ |
| $m$ | Migration rate | 2/365 |
| $s$ | Equation S6 | 6.00, 4.00, 4.00 |
| $f$ | Equation S6 | 2/365 |
| $\psi$ | Equation S6 | 1.25, 1.25, 2.00 |
| $g$ | Equation S7 | 0.0030, 0.0044, 0.0089 |
| $\mu$ | Table 1 | 1/730, 1/1095, 1/1825 |
| $L$ | Equation S4 | 1095, 730, 365 |
| $k$ | Table 1, Equation S4 | $(L^{-1} - \mu)N_0^{-1}$ |
| $\beta$ | Table 1 | $6 \times 10^{-6}$ |
| $\gamma$ | Table 1 | 1/10 |
| $\omega_A$ | Table 1 | 1/90 |
| $\omega_T$ | Table 1 | 1/1095, 1/730, 1/365 |

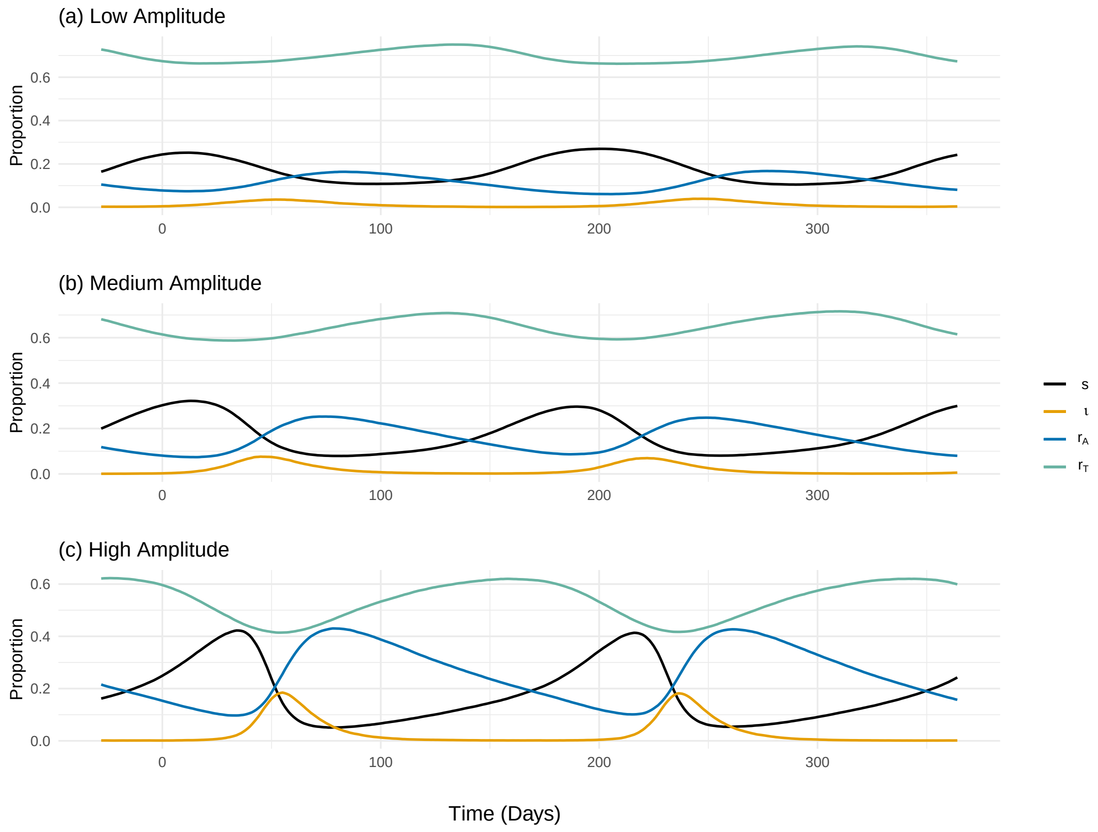


**Figure S2:** Examples of epidemiological dynamics in simulated populations. The parameter values in Table S2 generated the different (a) low, (b) medium, and (c) high amplitude curves for each compartment in the ODE model (1). Counts were converted to proportions after the simulations were completed.

Once the 300 synthetic populations were created, we sampled each population using the 27 sampling schemes given in table (S3) to mimic variation in serosurveillance data. We used binomial sampling with $n = 20$ trials to generate proportions of seropositive animals captured on each sampling day.

Interpolation and model fitting methods were then applied to these sampling schemes to test the accuracy of $\hat{\iota}$.

**Table S3:** Sampling schemes to mimic serosurveillance field data from 100 replicate populations in each category. Amplitude (amp.) category is either low, medium, or high. Mean amp. is the average amplitude calculated for each of the 100 datasets in each amplitude category. Seroprevalence (seroprev.) and prevalence (prev.) are the average prevalences calculated for each of the 100 datasets in each amplitude category, respectively. $\mathcal{R}_0$ is calculated from the parameters in table S2 using equation S2. Design describes when the day of sampling occurs.

| Scheme | Amp. | Mean Amp. | Seroprev. | Prev. | $\mathcal{R}_0$ | Design |
|---|---|---|---|---|---|---|
| 1 | Low | 0.080 | 0.11 | 0.013 | 5.7 | Daily |
| 2 | Low | 0.080 | 0.11 | 0.013 | 5.7 | Weekly |
| 3 | Low | 0.080 | 0.11 | 0.013 | 5.7 | Bi-Weekly |
| 4 | Low | 0.080 | 0.11 | 0.013 | 5.7 | Monthly |
| 5 | Low | 0.080 | 0.11 | 0.013 | 5.7 | Bi-Monthly |
| 6 | Low | 0.080 | 0.11 | 0.013 | 5.7 | Even Days |
| 7 | Low | 0.080 | 0.11 | 0.013 | 5.7 | Random Days |
| 8 | Low | 0.080 | 0.11 | 0.013 | 5.7 | 3-Day Clusters |
| 9 | Low | 0.080 | 0.11 | 0.013 | 5.7 | Week Clusters |
| 10 | Med | 0.17 | 0.16 | 0.018 | 5.6 | Daily |
| 11 | Med | 0.17 | 0.16 | 0.018 | 5.6 | Weekly |
| 12 | Med | 0.17 | 0.16 | 0.018 | 5.6 | Bi-Weekly |
| 13 | Med | 0.17 | 0.16 | 0.018 | 5.6 | Monthly |
| 14 | Med | 0.17 | 0.16 | 0.018 | 5.6 | Bi-Monthly |
| 15 | Med | 0.17 | 0.16 | 0.018 | 5.6 | Even Days |
| 16 | Med | 0.17 | 0.16 | 0.018 | 5.6 | Random Days |
| 17 | Med | 0.17 | 0.16 | 0.018 | 5.6 | 3-Day Clusters |
| 18 | Med | 0.17 | 0.16 | 0.018 | 5.6 | Week Clusters |
| 19 | High | 0.33 | 0.25 | 0.031 | 5.5 | Daily |
| 20 | High | 0.33 | 0.25 | 0.031 | 5.5 | Weekly |
| 21 | High | 0.33 | 0.25 | 0.031 | 5.5 | Bi-Weekly |
| 22 | High | 0.33 | 0.25 | 0.031 | 5.5 | Monthly |
| 23 | High | 0.33 | 0.25 | 0.031 | 5.5 | Bi-Monthly |
| 24 | High | 0.33 | 0.25 | 0.031 | 5.5 | Even Days |
| 25 | High | 0.33 | 0.25 | 0.031 | 5.5 | Random Days |
| 26 | High | 0.33 | 0.25 | 0.031 | 5.5 | 3-Day Clusters |
| 27 | High | 0.33 | 0.25 | 0.031 | 5.5 | Week Clusters |

#### APPENDIX 5: SUPPLEMENT TO THE RESULTS

We show results for the accuracy and functionality of our method on simulated data and then apply our method to Ebolaviruses in two African bat populations. We applied interpolation to all 27 simulated sampling schemes (tables S3, S4). Table (S4) gives the accuracy of the predicted peak, $\hat{\iota}_{peak}$, by calculating the magnitude of distance it is away from the true population peak, $\iota_{peak}$. Model fitting was assumed to work well on sampling schemes 1-5, and was applied to only sampling schemes 6-27 (table S5). Table (S5) again shows the accuracy of the prediction and also the size of the 95% CI and the proportion the true population peak falls within the CI.

**Table S4:** Results on the accuracy of estimating $\hat{r}_A(t)$ with interpolation for all sampling schemes. The accuracy metric, $|\iota_{peak} - \hat{\iota}_{peak}|$, is the magnitude of distance between the true population infectivity and predicted peaks. Distances between peaks are given in days.

| Scheme | Amp. | Design | $\lvert\iota_{peak} - \hat{\iota}_{peak}\rvert$ |
|---|---|---|---|
| 1 | Low | Daily | 7.35 |
| 2 | Low | Weekly | 14.14 |
| 3 | Low | Bi-Weekly | 18.96 |
| 4 | Low | Monthly | 30.90 |
| 5 | Low | Bi-Monthly | 24.53 |
| 6 | Low | Even Days | 17.36 |
| 7 | Low | Random Days | 20.24 |
| 8 | Low | 3-Day Clusters | 19.72 |
| 9 | Low | Week Clusters | 32.85 |
| 10 | Med | Daily | 4.37 |
| 11 | Med | Weekly | 7.61 |
| 12 | Med | Bi-Weekly | 11.26 |
| 13 | Med | Monthly | 17.09 |
| 14 | Med | Bi-Monthly | 29.94 |
| 15 | Med | Even Days | 11.02 |
| 16 | Med | Random Days | 12.39 |
| 17 | Med | 3-Day Clusters | 17.75 |
| 18 | Med | Week Clusters | 30.36 |
| 19 | High | Daily | 4.94 |
| 20 | High | Weekly | 5.17 |
| 21 | High | Bi-Weekly | 6.88 |
| 22 | High | Monthly | 10.49 |
| 23 | High | Bi-Monthly | 9.90 |
| 24 | High | Even Days | 4.87 |
| 25 | High | Random Days | 8.66 |
| 26 | High | 3-Day Clusters | 14.10 |
| 27 | High | Week Clusters | 25.02 |

**Table S5:** Results on the accuracy of estimating $\hat{r}_A(t)$ by fitting $r^*_A(t)$. The accuracy metric, $|\iota_{peak} - \hat{\iota}_{peak}|$, is the magnitude of distance between the true population infectivity and predicted peaks. CI size is the length of the 95% CI and we show the proportion of the true peaks that fall within this interval for each sampling scheme tested. All metrics are given in days.

| Scheme | Amp. | Design | $\lvert\iota_{peak} - \hat{\iota}_{peak}\rvert$ | CI Size | Prop. $\iota_{peak}$ in CI |
|---|---|---|---|---|---|
| 6 | Low | Even Days | 11.54 | 48.29 | 0.90 |
| 7 | Low | Random Days | 10.91 | 57.40 | 0.94 |
| 8 | Low | 3-Day Clusters | 9.77 | 49.96 | 0.90 |
| 9 | Low | Week Clusters | 14.52 | 71.18 | 0.91 |
| 15 | Med | Even Days | 6.21 | 26.51 | 0.91 |
| 16 | Med | Random Days | 7.03 | 25.15 | 0.88 |
| 17 | Med | 3-Day Clusters | 7.50 | 28.47 | 0.91 |
| 18 | Med | Week Clusters | 10.30 | 42.09 | 0.87 |
| 24 | High | Even Days | 3.03 | 16.03 | 0.96 |
| 26 | High | Random Days | 4.13 | 17.33 | 0.93 |
| 26 | High | 3-Day Clusters | 5.36 | 17.70 | 0.87 |
| 27 | High | Week Clusters | 10.22 | 24.66 | 0.66 |

Predicting the epidemic cycles using model fitting relies on the assumption of seasonal births occurring in the reservoir population. To test this assumption for Ebola virus (EBOV; *Zaire ebolavirus*) in *E. helvum* and *H. monstrosus*, we fit data on the proportion lactating individuals in population to equation (S6) using the likelihood function in equation (5) with the methods described in appendix 3. Figures (S3) and (S5) demonstrate that *E. helvum* has one cycle per year and *H. monstrosus* has two cycles per year, repsectively. Thus, we fix $f = 1/52$ weeks to generate the curves in figure (S4) to obtain the distribution of predicted peaks (figure 5) for *E. helvum*, and $f = 2/52$ weeks to generate the curves in figure (S6) to obtain the distribution of predicted peaks (figure 6) for *H. monstrosus*.

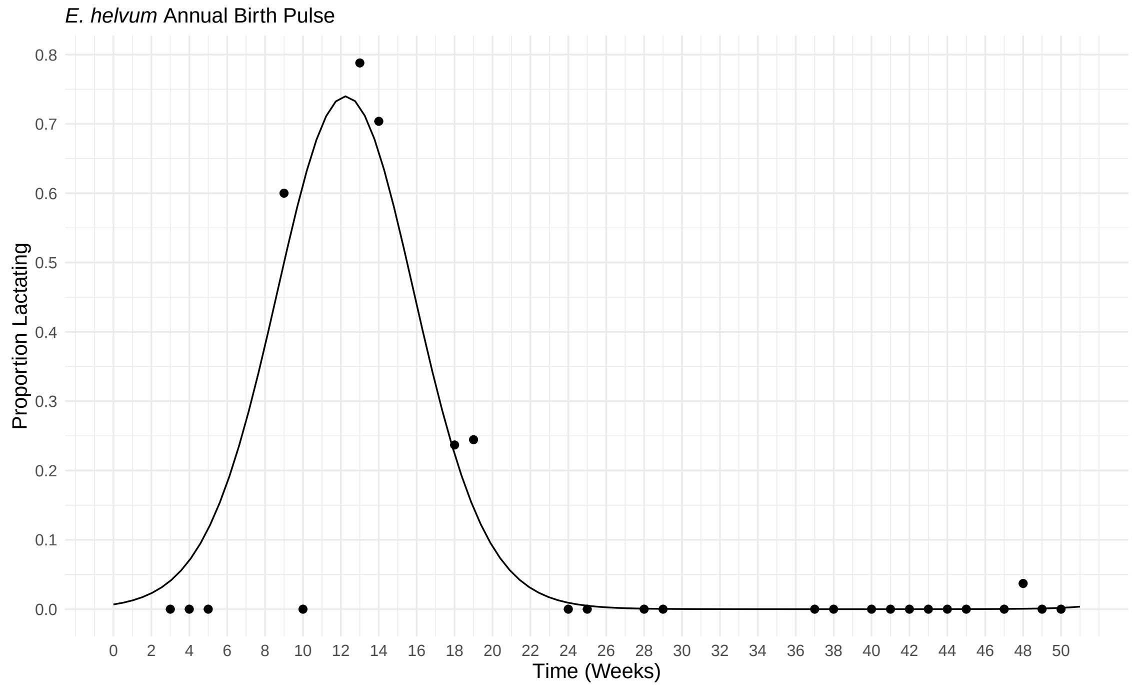


**Figure S3:** Estimated annual birth pulse in *E. helvum* from proportion lactating animals. Black dots represent the sampling data and the curve represents to estimated seasonal birth function.

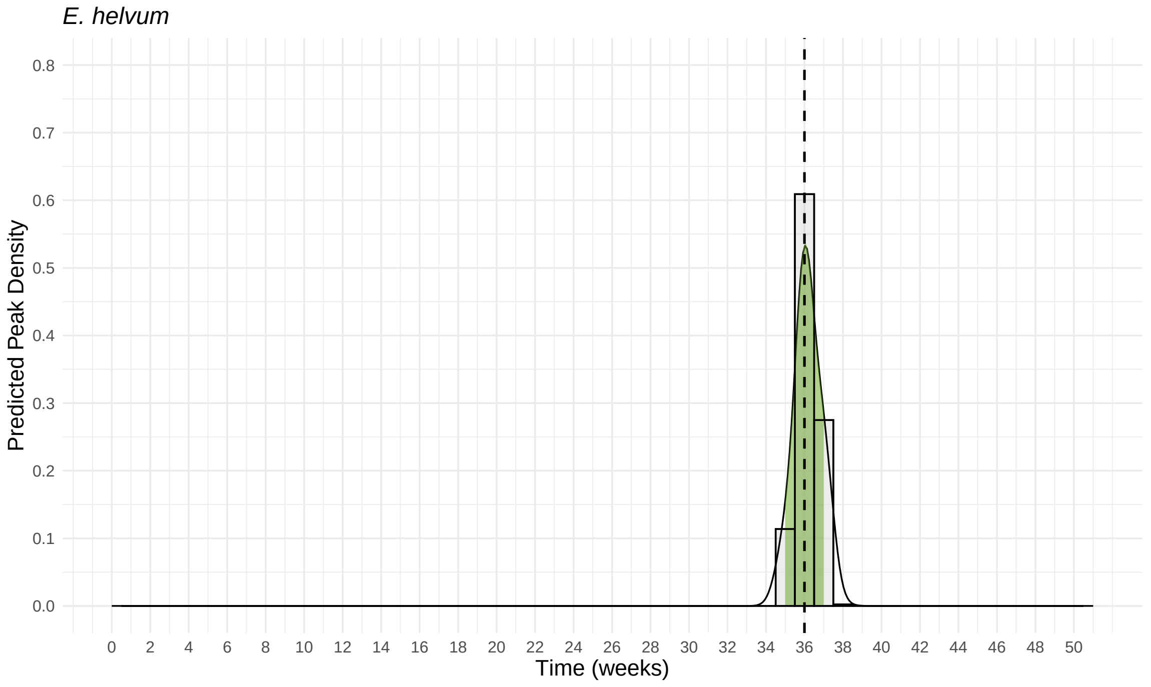


**Figure S4:** Distribution of weeks where the annual peak viral pulse was predicted for *E. helvum*. The vertical dashed line represents the mode and the green shaded area represents the 95% CI.

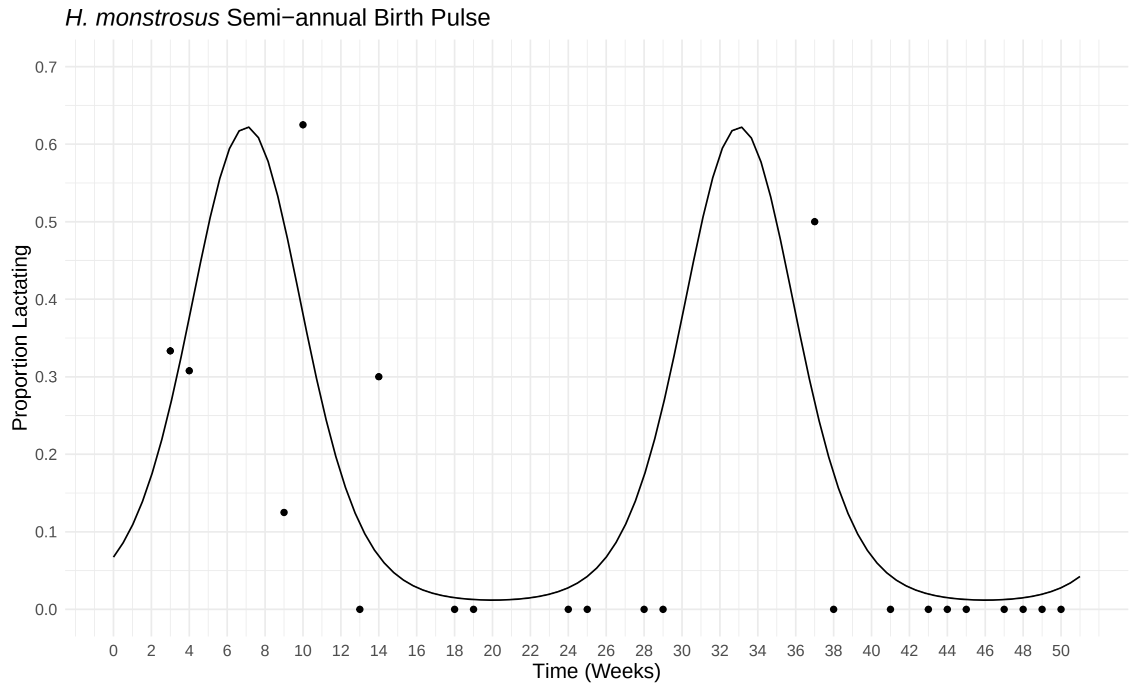


**Figure S5:** Estimated semi-annual birth pulse in *H. monstrosus* from proportion pregnant animals and annual precipitation patterns. Black dots represent the sampling data and the curve represents to estimated seasonal birth function.

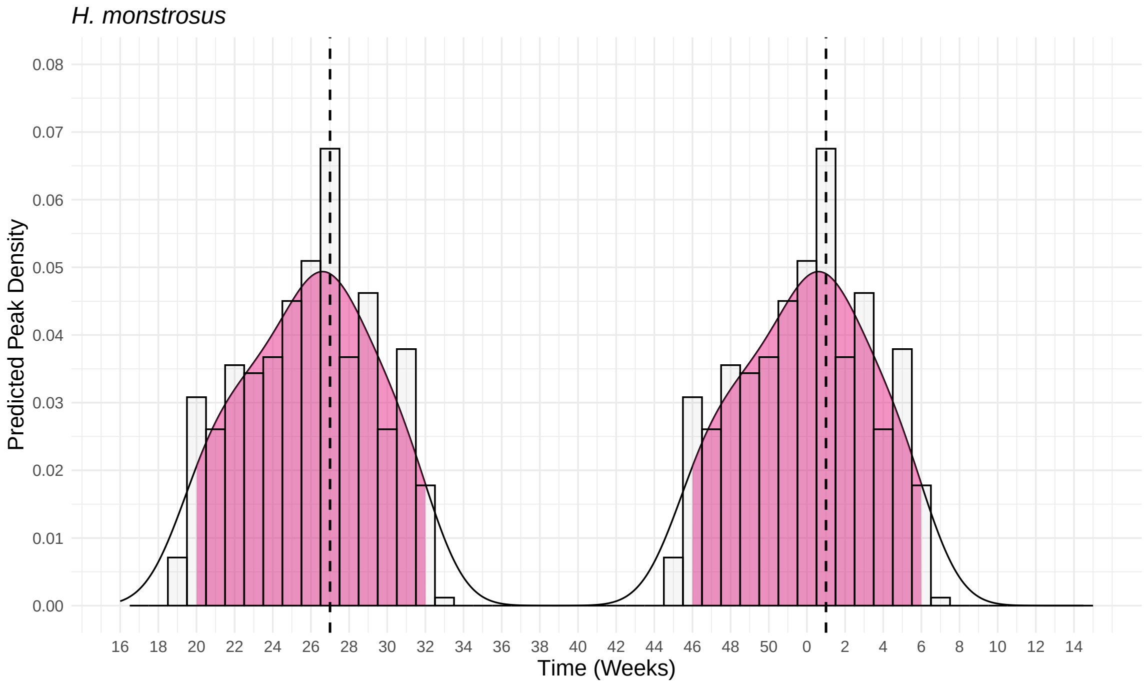


**Figure S6:** Distribution of weeks where the semi-annual peak viral pulses were predicted for *H. monstrosus*. The vertical dashed lines represent the modes and the pink shaded areas represent the 95% CIs for each pulse. Note the timescale on the x-axis begins at week 16 to accommodate the second peak that occurs at the end/beginning of each year.

#### APPENDIX 6: APPLICATION TO RANDOM EPISODIC SHEDDING

When surveillance data provides a good approximation for $\hat{r}_A$, interpolation can also be used to retrospectively understand episodic shedding in a reservoir population even in situations when episodic shedding does not coincide with seasonal birthing. For example, the prediction $\hat{\imath}(t)$ could be useful to understand the timing of spillover events in situations where a reservoir population routinely comes into contact with livestock or humans and causes disease. To demonstrate this additional utility of our interpolation method, we generate a reservoir population with a yearly seasonal birth pulse, but the dynamics of the circulating pathogen are such that pulses occur randomly with respect to reproduction. Parameter values for this simulation are given in table (S6), sampling from this population follows the protocol outlines in appendix 4, and the resulting curves are show in figure (S7).

**Table S6:** Parameter values used to generate random episodic shedding in a reservoir population. All rates are given in days.

| Parameter | Definition/Reference | Value in Simulations |
|---|---|---|
| $N_0$ | Initial pop. size | $1000$ |
| $m$ | Migration rate | 1/50 |
| $s$ | Equation S6 | 2.00 |
| $f$ | Equation S6 | 1/365 |
| $\psi$ | Equation S6 | 1 |
| $g$ | Equation S7 | 0.0030 |
| $\mu$ | Table 1 | 1/1095 |
| $L$ | Equation S4 | 730 |
| $k$ | Table 1, Equation S4 | $(L^{-1} - \mu)N_0^{-1}$ |
| $\beta$ | Table 1 | $8 \times 10^{-4}$ |
| $\gamma$ | Table 1 | 1/7 |
| $\omega_A$ | Table 1 | 1/70 |
| $\omega_T$ | Table 1 | 1/300 |

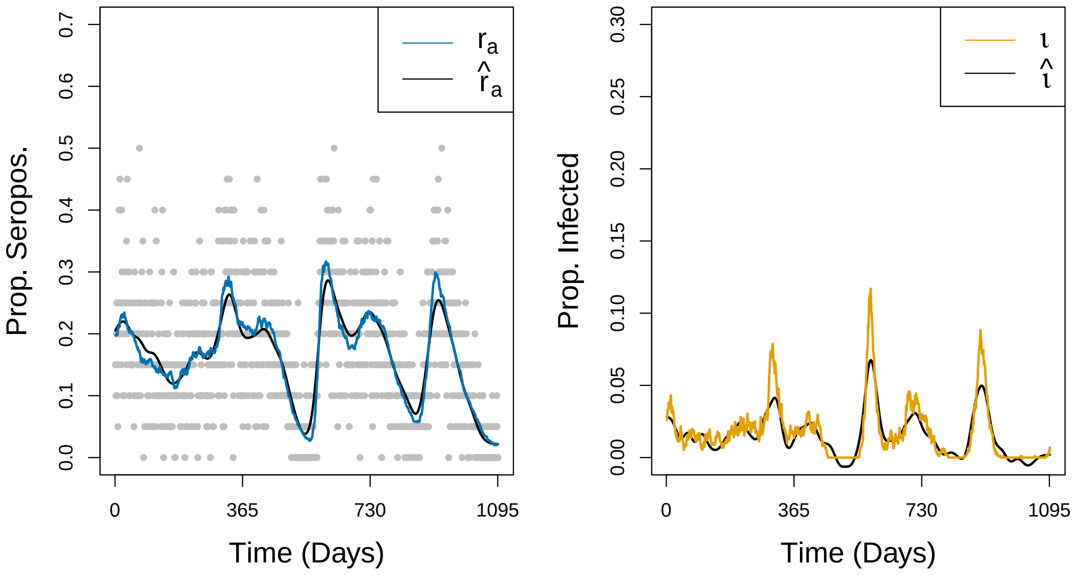


**Figure S7:** Results from using interpolation to daily sample serology data when episodic shedding occurs randomly. The grey points in the panels represent the raw simulated data.
